# Recommendations for tissue homogenisation and extraction in DNA metabarcoding of Malaise trap samples

**DOI:** 10.1101/2022.01.25.477667

**Authors:** Vera MA Zizka, Matthias F Geiger, Thomas Hörren, Ameli Kirse, Niklas W Noll, Livia Schäffler, Alice M Scherges, Martin Sorg

**Affiliations:** Leibniz Institute for the Analysis of Biodiversity Change (LIB), Zoological Research Museum Alexander Koenig (ZFMK), Centre for Biodiversity Monitoring and Conservation Science, Adenauerallee 160, 53113 Bonn, Germany; Entomological Society Krefeld (EVK), Marktstraße 159, 47798 Krefeld, Germany

**Keywords:** arthropod metabarcoding, biodiversity, monitoring, natura 2000

## Abstract

With increased application of DNA metabarcoding in fast and high-resolution biodiversity assessment, various laboratory protocols have been optimised in recent years and their further evaluation is subject of current research. Homogenisation of bulk samples and subsequent DNA extraction from destructed tissue is one way of starting the metabarcoding process. This essential step in the protocol can either be conducted from wet sample material (e.g. bulk insect samples) soaked in fixative or from completely dried individuals. While the latter method appears to produce more consistent results, it is time consuming and more prone to cross-contamination. We tested both homogenisation approaches with regard to time efficiency and biodiversity assessment of complex arthropod bulk samples, in particular how the amount of processed tissue affects taxon recovery. Both approaches reveal similar taxa compositions and detect a similar total OTU diversity in a single extraction reaction. Increased amounts of tissue used in DNA extraction improved OTU diversity detection and recovered particularly specific low-biomass taxa, making this approach valuable for samples with high biomass and/or diversity. Due to less handling time and lower vulnerability for cross-contamination we recommend the processing of wet material when sample homogenisation is applied.

## Introduction

For highly diverse groups as terrestrial arthropods and insects in particular, where morphological identification is difficult, slow and expensive, metabarcoding provides an efficient alternative (Bush et al., 2019; Evans et al., 2016; Morinière et al., 2019; van der Heyde et al., 2020; Yu et al., 2012). In recent years, a variety of studies evaluated and discussed promising sampling strategies (Gleason et al., 2020; Marquina et al., 2019; Pereira-da-Conceicoa et al., 2020; Steinke et al., 2020), laboratory procedures (Elbrecht et al., 2019; Majaneva et al., 2018; Piñol et al., 2019; Zizka et al., 2019), bioinformatic analyses (Boyer et al., 2016; Frøslev et al., 2017; Porter and Hajibabaei, 2020; Turon et al., 2020) and ways of integration into existing biodiversity monitoring matrices (Buchner et al., 2019; Cordier et al., 2018; Mächler et al., 2020; Pawlowski et al., 2018). This resulted in a variety of different DNA metabarcoding protocols, whereas standardisation is still lacking even though it is a major prerequisite for inter-comparability and transferability of methods to applied concepts (Bush et al., 2019; McGee et al., 2019; Pawlowski et al., 2018).

Aside from eDNA (environmental DNA) metabarcoding, where free extracellular DNA is processed (e.g. from soil, water, faeces), DNA can be extracted from enclosed communities (cDNA), more precisely, the sample’s fixative ethanol (Batovska et al., 2021; Hajibabaei et al., 2012; Martins et al., 2019; Zizka et al., 2019) or propylene glycol (Martoni et al., 2021), from added lysis buffer (Giebner et al., 2020; Ji et al., 2013; Kirse et al.,2021) or from homogenised tissue of specimens (Hardulak et al., 2020; Mata et al., 2020; Zizka et al., 2020). While the latter approach is currently considered most effective to assess biodiversity pattern (Hardulak et al., 2020; Marquina et al., 2019; Persaud et al., 2021; Zenker et al., 2020; Zizka et al., 2019) it prevents subsequent morphological determinations (Nielsen et al., 2019). Homogenisation and tissue-based DNA extraction can be conducted from wet (Beentjes et al., 2019; Gibson et al., 2015; Porter et al., 2019) samples in ethanol or from dried tissue after ethanol evaporation (Elbrecht et al., 2019; Hardulak et al., 2020; Hausmann et al., 2020; Steinke et al., 2020). While powder homogenate of dried samples usually appears finer than that of wet material, handling is more prone to cross contamination and time consuming. Since most DNA extraction approaches tolerate only a limited amount of tissue per reaction, only a subsample of complete material is usually processed, ranging between 1-100 mg (Elbrecht et al., 2017; Hausmann et al., 2020; Majaneva et al., 2018; Marquina et al., 2019; Mata et al., 2020). Higher tissue volume during DNA extraction requires multiple reactions or more voluminous DNA extraction kits and is increases effort and material costs. However, DNA extraction from the subsample tissue assumes perfect homogenisation and equal distribution within storage tubes, and it remains uncertain to what extent variations in tissue composition affects the assessment of species contained in bulk samples. Insight on that question is essential to decide on how to optimise diversity detection and is thus a prerequisite for successful and reliable application of tissue-based DNA metabarcoding. While Buchner et al. 2021 analyse the overlap in detected taxa between subsamples only for wet homogenisation of Malaise traps, other studies cover aquatic samples or other trapping types with lower biomass and diversity (Beentjes et al., 2019; Elbrecht et al., 2017; Mata et al., 2020). In addition, if several subsamples are applied, taxa detection is not compared amongst them or further methodological investigations are applied (e.g. comparison of different extraction methods (Majaneva et al., 2018). The overlap between subsamples of homogenised tissue or increases in taxon recovery by use of larger quantities of homogenised tissue in Malaise trap metabarcoding has never been tested systematically but bears important information for high resolution metabarcoding of terrestrial insect biodiversity.

Here we use five time-interval Malaise trap samples collected in a protected area in Germany and investigate the effect of homogenisation strategy and tissue subsampling on biodiversity assessments. Based on our results, we formulate best-practice recommendations for tissue-based DNA metabarcoding protocols esp. for Malaise trap samples, which ensure time and money efficiency, best quality of biodiversity assessment and also improved standardisation of DNA metabarcoding for biodiversity monitoring programs.

## Material and Methods

### Sampling

Samples were collected in the Nature reserve ‘Latumer Bruch’ near Krefeld in Western Germany. All samples originate from one Malaise trap (51.326701N, 6.632973E). Detailed information about samples taken between May and July are given in Table 1.

**Tab. 1:** Malaise trap samples analysed: duration: time collection bottle was installed on the Malaise trap, wet biomass: complete wet biomass (g) and biomass per day (g/day) over sampling interval

| ID | duration | wet biomass (g) | wet biomass (g/day) |
| --- | --- | --- | --- |
| T 1 | 12.05 - 18.05.19 | 14.1 | 2.4 |
| T 2 | 29.05 - 08.06.19 | 65.9 | 6.7 |
| T 3 | 28.06 - 07.07.19 | 71.7 | 8 |
| T 4 | 07.07 - 18.07.19 | 40 | 3.6 |
| T 5 | 18.07 - 28.07.19 | 70.1 | 7 |

Malaise trap sampling was conducted in a standardised manner (for details see Ssymank et al., 2018). Samples were collected in 96% denatured ethanol (1% MEK). After collection, ethanol was replaced with new 96% undenatured ethanol and stored at −20°C for further processing.

### Laboratory work

Supernatant ethanol was removed and each sample was separated into two size classes by sieving of wet specimen through a 4 mm x 4 mm mesh with a wire diameter of 0.5 mm (untreated stainless steel). In the following, the size fractions will be referred to as either S (small, < 4mm) or L (large, > 4 mm). Depending on sample volume, individuals of both size classes were transferred to 30 ml tubes (Nalgene, wide-mouth bottle, polypropylene) or to 50 ml Falcon tubes, and approximately 20 g of grinding balls (5 mm diameter, stainless steel, Retsch GmbH) were added to each sample. Homogenisation of wet samples was conducted with a mixer mill MM 400 (Retsch GmbH) at a frequency of 30 s^-1^ for five minutes. Nine subsamples per sample and size class were transferred to 1.5 ml Eppendorf tubes, centrifuged for 1 minute at 10,000 rpm (Heraeus Fresco 21 Centrifuge, Thermo Scientific) and dried overnight in a shaking incubator (ILS6, VWR) at 50 °C (90 samples in total, 5 samples x 2 treatments x 9 subsamples). Subsamples were weighed (20 mg ± 6 mg) on a fine scale balance (Entris, Sartorius). Together with 6 negative controls (200 μl lysis buffer, no sample added) and DNA was extracted with the DNeasy 96 Blood and Tissue Kit (Quiagen, Hilden, Germany) following manufacturer instructions. Remaining sample tissue (in 30 ml tubes, not transferred to 1.5 Eppendorf tubes) was centrifuged for 1 minute at 4700 rpm (MegaStar 1.6, VWR Collection) and the supernatant discarded afterwards. Remaining tissue was left to dry in a shaking incubator at 50 °C for up to 3 days until complete ethanol evaporation. Dried tissue of samples containing less than 30 ml was again homogenised at a frequency of 30 s/sec for 5 minutes with the Retsch mixer mill (MM400). Samples consisting of more than 30 ml source material were homogenised for 3 minutes with a Turax mixer mill (Tube Mill 100 Control) at 25,000 rpm because dried material was clustered to a hard unit and could not be destructed with the former mill. Nine subsamples per sample were transferred to 1.5 Eppendorf tubes and weighed (23 mg ± 5 mg). To ensure processing of identical samples during this experiment, samples already homogenised under wet condition were dried and homogenised again under dry condition. Obviously, this includes an additional homogenisation step for those samples, which likely influences fineness of material. This will be picked up in the discussion. For simplicity, the former approach will hereinafter will be referred to as *wet homogenisation*, while the latter approach (wet with additional dry homogenisation) will be referred to as *dry homogenisation*. Together with 6 negative controls DNA was extracted with the DNeasy 96 Blood and Tissue Kit (Quiagen, Hilden, Germany) following manufacturer instructions for both plates. Extraction success and DNA quality was checked on a 1 % agarose gel.

A two-step PCR protocol was applied using standard Illumina Nextera primers for dual indexing of samples. The first PCR was performed with the PCR Multiplex Plus Kit (Qiagen, Hilden, Germany) using 12.5 μl master mix, 1 μl of DNA template, 0.2 μM of the fwhF2 forward (Vamos et al., 2017) and Fol_degen_rev reverse (Yu et al., 2012) primers respectively. The primer pair targets a 313 bp long stretch of the COI DNA barcode region and was positively evaluated in a comparison on Malaise trap samples (Elbrecht et al., 2019). The PCR mix was filled up with 10.5 μl ddH_2_O to a 25 μl reaction volume. The following PCR program was applied: initial denaturation at 95 °C for 5 min; 25 cycles of: 30 s at 95 °C, 30 s at 50 °C and 50 s at 72 °C; final extension of 5 min at 72 °C. First step PCR (PCR 1) product was used for the second PCR (PCR 2), also conducted with the PCR Multiplex Plus Kit (Qiagen, Hilden, Germany). Reaction included 1 μl DNA template from PCR 1, 0.2 μM of each tagging primer (Nextera, Illumina, San Diego, USA) and 12.5 μl master mix filled up with 10.5 μl H_2_O. Primers included a nucleotide overhang as a binding site for primers in PCR 2, which was run with the same program as PCR 1 but with only 15 cycles. PCR success was evaluated on a 1 % agarose gel before PCR products were normalised using a SequalPrep Normalisation plate (Thermo Fisher Scientific, MA, USA) following the manufacturer’s instructions with an end concentration of 25 ng per sample (100 μl). Of each sample 10 μl was pooled together and a left sided size selection was applied twice on the sample pool to remove primer residuals (ration 0.76x, SPRIselect Beckman Coulter). Library concentration was measured with a Quantus fluorometer (Promega, Madison, USA) and on a fragment analyser (Agilent Technologies, Santa Clara, CA, USA) and the pool was sent for sequencing on two Miseq runs (2 × 300 bp) to Macrogen Europe B.V., Netherlands.

## Data analysis

The quality of sequences delivered by Macrogen was determined through the program Fastqc (Andrews et al., 2012). Subsequent data processing was conducted for all samples as implemented in JAMP v0.67 (https://github.com/VascoElbrecht/JAMP) using standard settings. Paired-end reads were merged (Edgar and Flyvbjerg, 2015) (module Merge_PE) with vsearch v2.15.0 (Rognes et al., 2016). Cutadapt 3.4 (Martin, 2017) was used to remove primers and to discard sequences of unexpected length so that only reads with a length of 303 - 323 bp were used for further analyses. The module Max_ee was used to discard all reads with an expected error >0.5 (Edgar and Flyvbjerg, 2015). Sequences were dereplicated, singletons were removed and sequences with ≥97% similarity were clustered into Operational Taxonomic Units (OTUs) using uparse (Cluster_otus) (Edgar, 2013). OTUs with a minimal read abundance of 0.003% per sample were retained for further analysis and the program LULU was used for further qualitative filtering (Frøslev et al., 2017). Taxonomic assignment of molecular units was conducted by comparison with an Arthropoda reference database. The database was created by taxalogue (https://github.com/nwnoll/taxaloguecommit:62ce71819af40a6e605e9142f0ccd69318477596) with sequences from BOLD (https://www.boldsystems.org), NCBI GenBank (https://www.ncbi.nlm.nih.gov/genbank/) and GBOL (https://bolgermany.de/gbol1/ergebnisse/results). Taxon names were normalised according to the NCBI Taxonomy (https://www.ncbi.nlm.nih.gov/taxonomy). Only those OTUs were kept that had at least 85% similarity with a sequence in a reference database (using vsearch version 2.14.1 (Rognes et al., 2016) with –usearch_global command (Edgar, 2010) and the following parameters: –id 0.85, –dbmask none, –qmask none, –maxhits 1000, blast6out and –maxaccepts 0). Custom scripts were used to extract the best hits and assign each OTU a taxonomic name. For analysis of significant differences in OTU numbers between homogenisation methods and size fractions, a Kruskal-Wallis test was performed as integrated in the R package “dplyr” (Wickham et al., 2021). Dissimilarity indices were calculated with the package “vegan” (Oksanen et al., 2019). For species accumulation curves and associated calculations of extrapolated values the package “iNEXT” (Hsieh et al., 2016) was used. Figures were constructed with the R package ggplot (Wickham, 2016).

## Results

On the two Miseq runs 13.3 and 14.7 million reads in forward and reverse direction were assigned to the specified index combinations. Raw data are uploaded at GenBank (accession number: XX). After quality filtering, on average 114,394 (± 20,365) reads were kept per sample. For both homogenisation approaches combined and all subsamples, in total 1,529 OTUs were clustered with the following order-level assignments: Coleoptera 158 (10.3%), Diptera 372 (24.3%), Hemiptera 134 (8.8%), Hymenoptera 689 (45.1%), Lepidoptera 52 (3.4%), other orders 99 (6.4%), no assignment to order level 25 (1.6%). Of those, 1088 OTUs were assigned to genus or species level. Detailed information about OTU assignments and read distribution on order level are summarised in Table 1 and Table S1. While the highest number of OTUs was assigned on average to dipterans (L: 41.6%, S: 37.5%) and hymenopterans (L: 38%, S: 32.3%), the main proportion of reads was related to dipterans (L: 60.8%, S: 77%) and < 10% to representatives of Hymenoptera (Table 1). This was most pronounced for size fraction S, where on average only 4.7% of the reads were assigned to this highly diverse order.

**Table 1:** Average proportion of OTUs and reads assigned per time interval and through all samples to the orders Coleoptera, Diptera, Hemiptera, Hymenoptera, Lepidoptera, other arthropod orders and OTUs not assigned to order level (see table S1) for both homogenization approaches and all subsamples combined

| Order | Size | OTU number [%] | Read number [%] | Total OTU number |
| --- | --- | --- | --- | --- |
| Coleoptera | L | 5.7 ± 1.9 | 6 ± 4.9 | 158 (10.3%) |
| | S | 11.1 ± 1 | 5.5 ± 2.8 | |
| **Diptera** | **L** | **41.6 ± 9.8** | **60.8 ± 13.3** | **372 (24.4%)** |
|  | **S** | **37.5 ± 8.8** | **77 ± 11.9** |  |
| Hemiptera | L | 3 ± 0.8 | 0.4 ± 0.3 | 134 (8.8%) |
| | S | 11.2 ± 2.3 | 10.1 ± 10.7 | |
| Hymenoptera | L | $38 \pm 5.2$ | $18.7 \pm 7.9$ | 689 (45.1%) |
| | S | $32.3 \pm 7.5$ | $4.7 \pm 2.5$ | |
| Lepidoptera | L | $7.6 \pm 3.2$ | $13.8 \pm 5.4$ | 52 (3.4%) |
| | S | $2.9 \pm 1.3$ | $1.8 \pm 1$ | |
| Other arthropods | L | $2.8 \pm 0.8$ | $0.3 \pm 0.2$ | 98 (6.4%) |
| | S | $3.6 \pm 0.6$ | $0.2 \pm 0.1$ | |
| No assignment | L | $1.2 \pm 0.4$ | $0.1 \pm 0.2$ | 25 (1.64%) |
| | S | $1.3 \pm 0.3$ | $0.7 \pm 0.5$ | |

While different emptying dates and the different size classes per sample showed distinct community compositions (Fig. 2, p < 0.002), the condition during homogenisation (only wet or wet and additional dry homogenisation) had no effect on sample ordination in NMDS analysis (p = 0.997, Fig. 2). However, average Jaccard dissimilarity between subsamples homogenised in wet and additional dry condition (0.179 ± 0.06) was lower (p < 0.001) than dissimilarities between subsamples homogenised in only wet condition (0.207 ± 0.081) and when subsamples of one emptying date and size fraction were compared amongst homogenisation approaches (0.206 ± 0.089, p < 0.001). Bray-Curtis dissimilarity was on average lower within wet and dry (0.11 ± 0.07) compared to only wet (0.11 ± 0.09) homogenised subsamples than comparing subsamples between methods (0.13 ± 0.08, p < 0.01).

**Figure 1:**
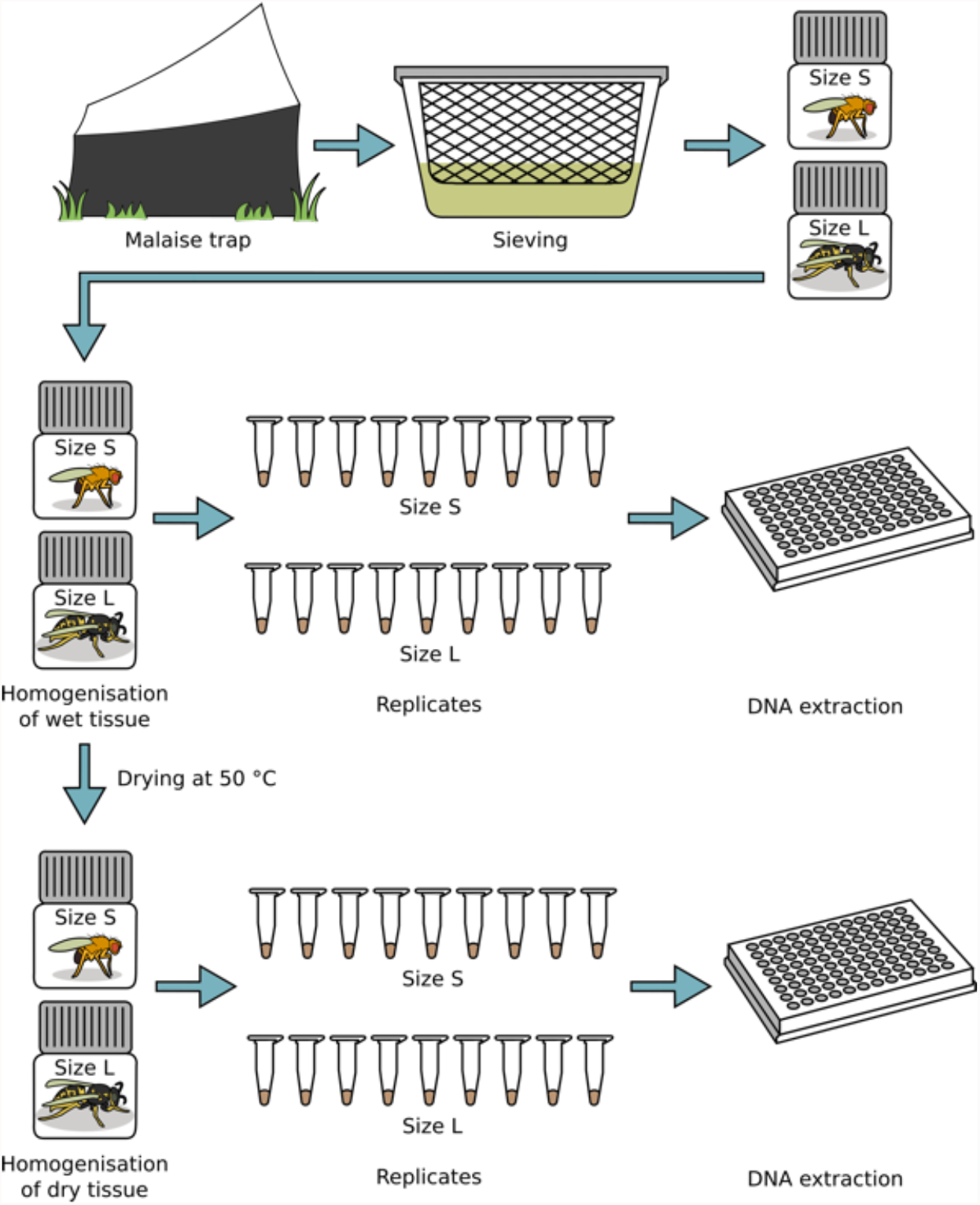
Experimental setup. Malaise trap samples were sieved in ethanol and separated into size fraction **L**arge (>4mm) and **S**mall (<4mm). Wet tissue was homogenised and nine subsamples per size fraction with ∼20 mg each were transferred to 1.5 μl Eppendorf tubes for separate DNA extraction. Homogenised tissue was dried and again homogenised in dry conditions. Again, nine subsamples per size fraction with ∼20 mg each were transferred to 1.5 μl Eppendorf tubes. Subsequent DNA workflow was conducted as described in Material and Methods.

**Figure 2:**
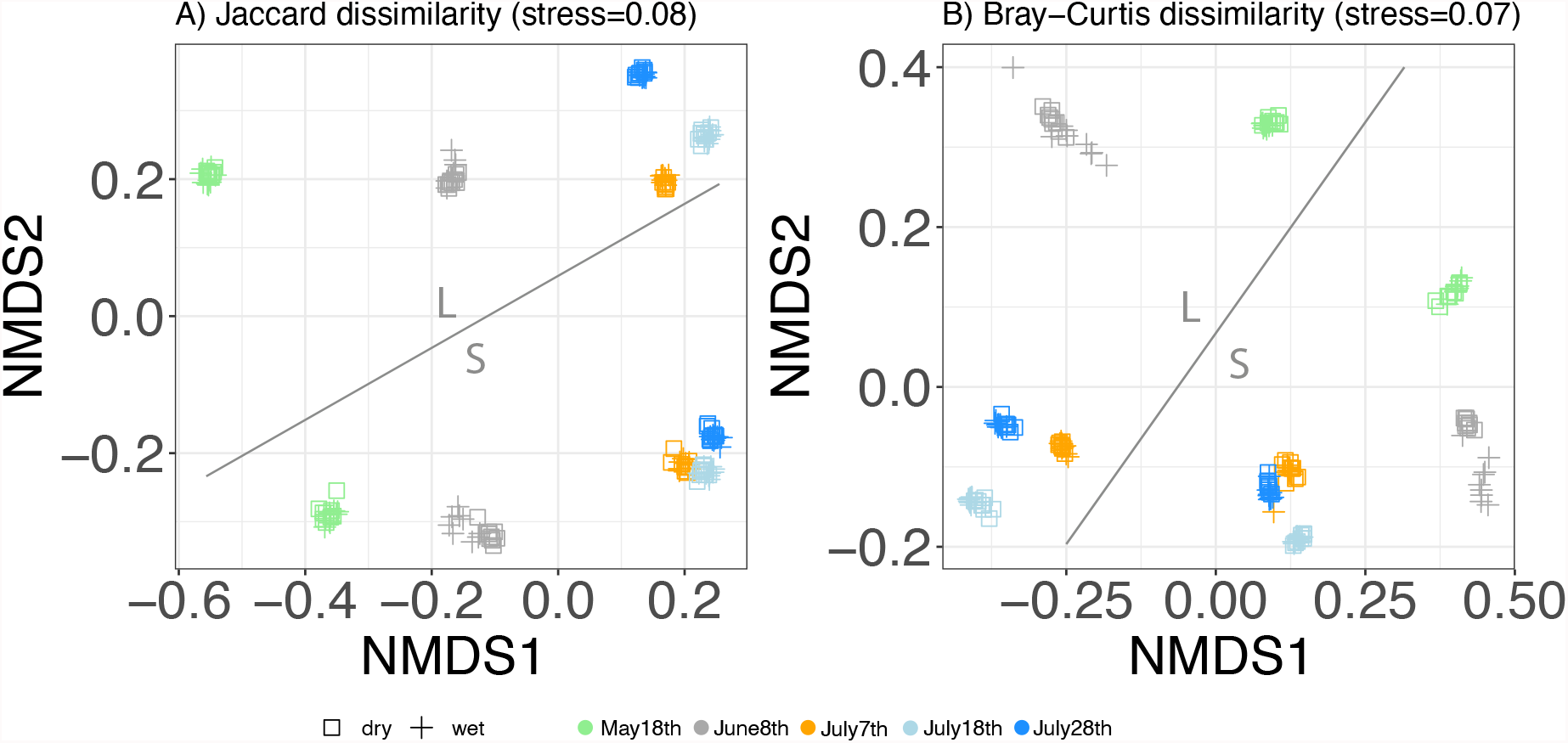
Non-metric multidimensional scaling based on A) Jaccard (presence/absence data) and B) Bray-Curtis (abundance data – regarding read numbers) dissimilarity matrices. Samples include the nine subsamples per emptying date (colour coding), size fraction (S and L, marked in figure) and condition during homogenisation (shape coding).

For samples processed under wet condition, on average 60.7% ± 8.5 of calculated total diversity could be detected in size fraction S and 75.8% ± 7.9 in size fraction L when only one subsample was processed in extraction (∼20 mg of tissue, Fig. 3). With nine extraction subsamples (9 x ∼20 mg) 88.4% ± 3.2 and 92.4% ± 4.2 of calculated total richness were detected in fraction S and L respectively, while 95% of calculated species richness was assessed with 12 ± 5.4 and 17 ± 3.4 subsamples. For samples homogenised in wet and additional dry condition, a single extraction (∼20 mg) of size class S revealed 64% ± 2.3 of calculated species richness, while 79% ± 3.8 could be detected in 20 mg of size class L (Fig. 3). On average 88.9% ± 1.6 of total diversity was assessed with the nine applied subsamples for size fraction S and 93.1% ± 3.8 for size fraction L, while 95% of total was calculated with 16 ± 2.1 (S, 320 mg) and 13 ± 7.1 (L, 260 mg) subsamples (∼320 mg and ∼260 mg, Fig. 3).

**Figure 3:**
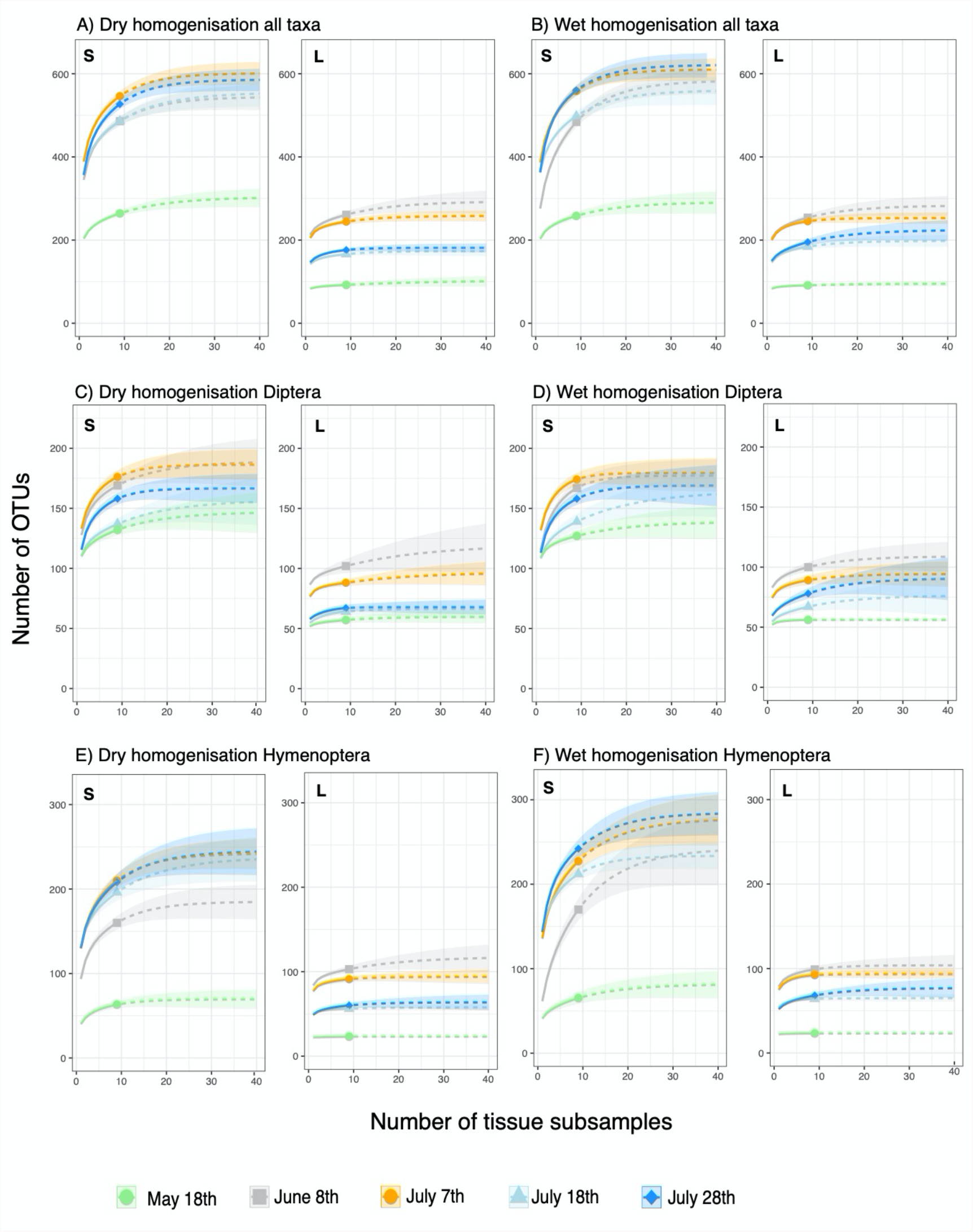
Species accumulation curves. Increased number of OTUs with increased amount of processed tissue in extraction is illustrated as well as extrapolation up to 40 subsamples. All taxa homogenized in A) dry and B) wet condition; Diptera recovered in samples homogenised in C) dry and D) wet condition; Hymenoptera detected in samples homogenised in E) dry and F) wet condition. Size fraction is marked in the upper left corner of each subfigure (S = small, L = large).

**Figure 4:**
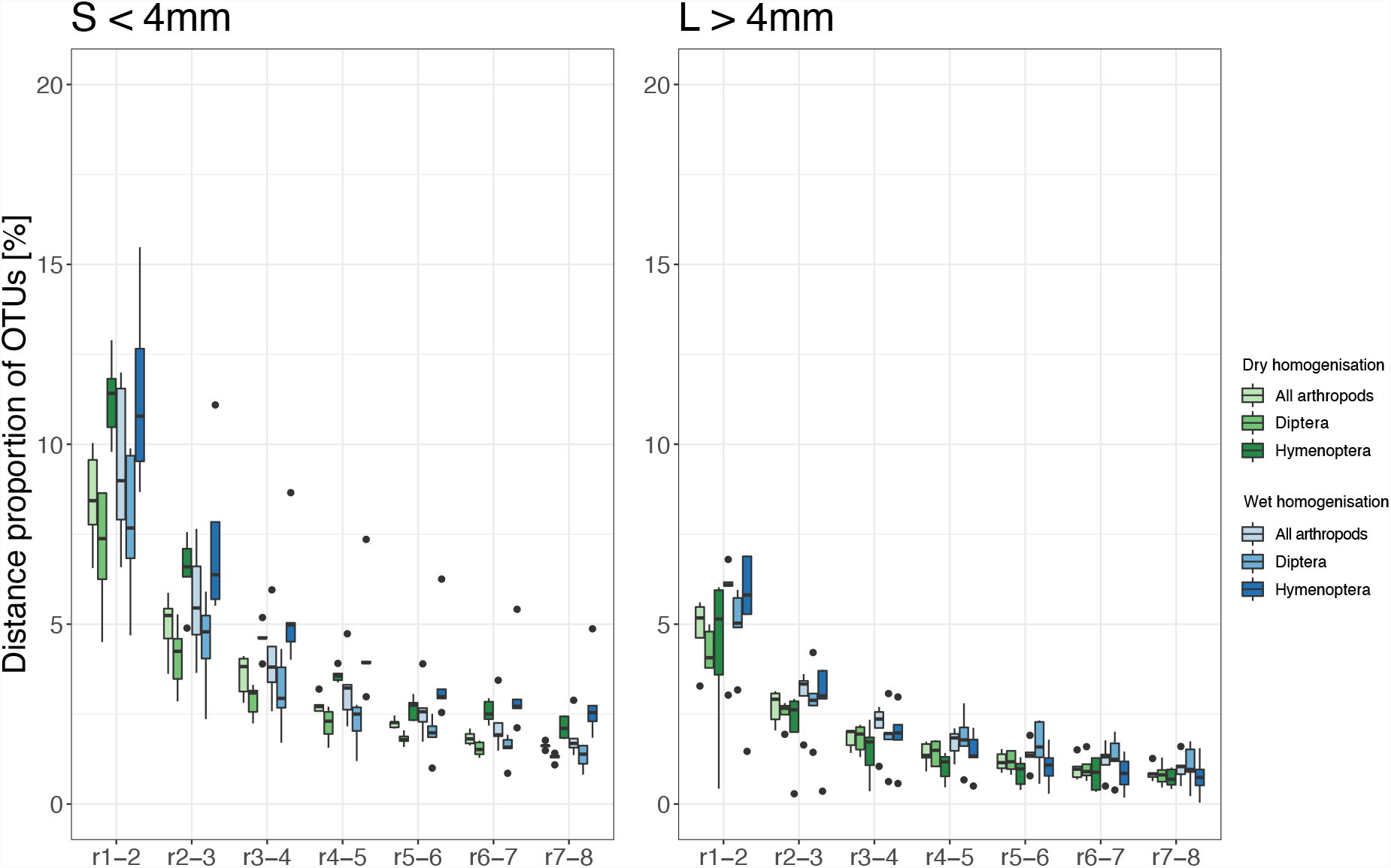
Proportion of molecular units (OTUs) detected with additional DNA extraction subsamples. A) for samples of size fraction L (> 4mm) and B) for size fraction S (< 4 mm). The x-axis describes differences between processed subsamples (e.g. r1-r2 difference in relative number of OTUs detected with one subsamples (∼20 mg) compared to two subsamples (∼40 mg)).

Detailed analysis of Hymenoptera in only wet homogenised tissue revealed on average 47.2% ± 13.3 of extrapolated total species richness in size fraction S and 80.4% ± 10.4 in size fraction L when a single subsample was processed (Fig.3). Increased to nine subsamples, 81.4% ± 7.6 (S) and 96.2% ± 4.7 (L) of calculated total diversity were detected. Extrapolations revealed, that 95% of total calculated diversity was achieved by processing 22 ± 5.2 (S, ∼440 mg) and 7.4 ± 6 (L, ∼140 mg) replicates. For wet and additional dry homogenisation, one extraction subsample revealed 54% ± 2.7 (S) and 82.4% ± 9.6 (L) of total species richness, while 86.2% ± 3.1 and 95.1% ± 4.8 could be assessed with nine extraction subsamples. Calculations revealed a detection of 95% from total hymenopteran species richness if 18 ± 4 (360 mg) and 9.8 ± 8 (200 mg) replicates were processed (Fig. 3).

For wet homogenisation, in detail analysis of dipteran representatives revealed 70.6% ± 5.6 of calculated diversity in size fraction S and 77% ± 10.3 in size fraction L, when only a single extraction subsample was processed. Detected richness increased to 92.1% ± 4.7 and 91% ± 5.6 when nine subsamples were processed (95% were reached with 14.2 ± 7.9 and 5 ± 1.9 subsamples). With additional dry homogenisation processing of one tissue subsample revealed 70.6% ± 2.7 of extrapolated diversity for size fraction S and 81.1 ± 5.9 for size fraction L. With nine extraction subsamples, taxa detection increased to 91.2% ± 3.4 (S) and 93.3% ± 5.5 (L). Calculations revealed a detection of 95% from total Diptera if 15.6 ± 5.4 (∼320 mg) and 14.2 ± 10.2 (∼200 mg) replicates were processed. For detailed information about observed and calculated species richness see Figure 3.

## Discussion

Homogenisation of bulk samples and subsequent DNA extraction from destructed tissue is widely applied when insect biodiversity is assessed through DNA metabarcoding (Beermann et al., 2021; Hardulak et al., 2020; Mata et al., 2020). It is up to now more efficient for bulk sample analysis than other extraction methods (Marquina et al., 2019; Persaud et al., 2021; Zenker et al., 2020). Here we set out to test different homogenisation protocols and how subsampling of homogenised tissue affects diversity estimates of highly diverse Malaise trap samples.

### Comparison of different homogenisation approaches

The average dissimilarity between reactions homogenised under dry conditions was lower than dissimilarity between reactions processed under wet condition for presence/absence analysis, which indicates a higher homogeneity of tissue samples processed under dry condition. To ensure comparability and processing of identical sampling material (as stated in Material and Methods section line 150-156), the dry homogenisation approach was based on material already destructed under wet condition. This additional step of 3 min homogenisation most probably influenced refinement of samples and similarity of subsamples. In addition, on average a lower tissue weight was processed subsequent to wet homogenisation due to a weight decrease after drying (difference 3 mg). However, Jaccard’s dissimilarity indices of subsamples which were homogenised in wet condition was on average 0.21 ± 0.08, mainly due to high inconsistencies between subsamples from June 8^th^ with an average dissimilarity of 0.35 ± 0.02 compared to dissimilarities between subsamples of the other collection dates (0.18 ± 0.06; Fig. 2). The homogenisation of dried samples implements drying for approximately 48 h at temperatures around 50 °C to guarantee the complete evaporation of ethanol from the sample. In addition, the fine powder resulting from dry homogenisation is electrostatically charged and thus bears a high risk of cross contamination between samples and general lab contamination, and further increases the time for sample handling (Buchner et al., 2021; Elbrecht and Steinke, 2019). In comparison, the homogenisation of wet samples soaked in ethanol circumvents the drying step and is therefore more time efficient and suitable for large scale approaches as implemented in several studies on aquatic samples (Hajibabaei et al., 2019; Majaneva et al., 2018; Pereira-da-Conceicoa et al., 2020). In addition, the handling of homogenised tissue in ethanol is simplified and reduces contamination risk (Elbrecht and Steinke, 2019). The minor differences we observed between the two applied methods and the above-mentioned experimental setup allows to recommend homogenisation of wet material for tissue-based DNA metabarcoding Malaise trap samples as it reduces processing time and contamination risk. Additionally, results indicate, that an additional homogenisation step of dried material as e.g. through bead-grinding should be integrated after material has been subsampled to increase the fineness of material as also conducted in Buchner et al., 2021.

### Amount of tissue needed for homogenisation

A high, but incomplete proportion of the total insect diversity was assessed when processing a single tissue subsample in extraction, which opens up an additional perspective to previous studies highlighting wet homogenisation of samples (Buchner et al., 2021). This accounts for both tested variables, basis material for homogenisation (wet or dry) and size fractions (Fig. 3). For all samples at least 60% of calculated total arthropod diversity were detected with a single extraction from ∼20 mg of tissue (Fig. 3 A+B), revealing strong alterations between size fractions time intervals and higher insect taxa. The processing of additional subsamples only moderately increase diversity estimates for size fraction L and samples of both size fractions from samples collected in May. While those samples constitute a comparatively low biodiversity, a strong increase was detected for remaining samples of the small size fractions and collected in summer. Results indicate that extraction should be conducted from more tissue for highly diverse samples, either through increasing number of extraction subsamples, as it was also recommended in (Elbrecht and Steinke, 2019) or a higher tissue volume per DNA extraction. Additional rare material for extraction however comes with higher expenses but can increases taxa detection by 25-30% as an average over all taxa. Referring to the 3000 BINs detected within a year-long Malaise trap collection through single-specimen barcoding in Geiger et al., 2016 this can result in 600-900 additional molecular units for year-long application of a malaise trap.

The most pronounced increase of OTUs with additional extraction subsamples was observed for representatives of the order Hymenoptera, which is almost twice as high as for Diptera (Fig. 3, Tab. 2). Representatives of these two orders are the main targets in many Malaise trapping studies (Ssymank et al., 2018). However, as also indicated with previous studies, dipterans are present in much higher individual numbers, constituting a higher proportion of biomass in Malaise trap catches in Germany, while diversity of both groups are considered to be similar (Geiger et al., 2016). An underrepresentation of specific insect families, especially constituting taxa of low-biomass (Elbrecht et al., 2021) has been reported from previous metabarcoding studies (Elbrecht and Steinke, 2019; Krehenwinkel et al., 2017; Yu et al., 2012). This accounts e.g. for highly diverse parasitoid hymenopterans, which depict important ecosystem functions. While insufficient primer binding efficiency is discussed as the main reason for this phenomenon, our results indicate, that the extraction from an insufficient amount of raw tissue material could also bias detected diversity pattern. It also assumes that taxon biomass in complex bulk samples is affecting detection probability if a limited amount of tissue is processed in extraction but can be increased through insertion of higher tissue amounts. Again, this accounts especially for highly diverse samples as indicated here through the small size fraction. We demonstrate, that the use of higher amount of tissue of approx. 180 mg over nine subsamples increase the detection of low biomass taxa, even if general sequence coverage remains low (5% of reads assigned to Hymenoptera, Table 1). Additionally, the application of higher sequencing depth can increase taxa recovery and overlap between extraction subsamples. We here only tested a sequencing depth of on average 114,394 ± 20,365 reads per sample (after quality filtering) and detailed analysis to understand the linkage between higher sequencing depth and replication strategy is out of the scope of the present study. Further investigation could reveal an increase in sequencing depth as the most effective way to optimise taxon recovery under financial constraints.

## Conclusion

We recommend homogenisation of wet material in tissue-based DNA metabarcoding of Malaise trap samples due to similar levels of taxon recovery from dry and wet tissue homogenisation combined with a lower time effort and contamination susceptibility of the latter approach. In both, dry and wet homogenisation, additional DNA extractions from more tissue results in a higher number of detected taxa, in particular those of low biomass. The amount of processed tissue and number of subsamples affects the resolution taxa-specifically. A decision on the more complete diversity detection associated with higher resources depends on the focused scientific goals of individual studies.

## Supporting information

OTU table with taxonomic assignment and plausibility check for species level identification

## Data Accessibility

The raw sequence used in this study is available at NCBI SRA: XX

## Acknowledgements

This work was supported by the Ministry for Environment, Agriculture, Conservation and Consumer Protection of the German State of North Rhine-Westphalia (No. III-1-620.08)

